# Spike sorting AI agent

**DOI:** 10.1101/2025.02.11.637754

**Authors:** Zuwan Lin, Arnau Marin-Llobet, Jongmin Baek, Yichun He, Jaeyong Lee, Wenbo Wang, Xinhe Zhang, Ariel J. Lee, Ningyue Liang, Jin Du, Jie Ding, Na Li, Jia Liu

## Abstract

Spike sorting is a fundamental process for decoding neural activity, involving preprocessing, spike detection, feature extraction, clustering, and validation. However, conventional spike sorting methods are highly fragmented, labor-intensive, and heavily reliant on expert manual curation, limiting their scalability and reproducibility. This challenge has become more pressing with advances in neural recording technology, such as high-density Neuropixels for large-scale neural recording or flexible electrodes for long-term stable recording over months to years. The volume and complexity of these datasets make manual curation infeasible, requiring an automated and scalable solution. Here, we introduce SpikeAgent, a multimodal large language model (LLM)-based AI agent that automates and standardizes the entire spike sorting pipeline. Unlike traditional approaches, SpikeAgent integrates multiple LLM backends, coding functions, and established algorithms, autonomously performing spike sorting with reasoning-based decision-making and real-time interaction with intermediate results. It generates interpretable reports, providing transparent justifications for each sorting decision, enhancing transparency and reliability. We benchmarked SpikeAgent against human experts across various neural recording technology, demonstrating its versatility and ability to achieve curation consistency that are equal to, or even higher than human experts. It also drastically reduces the expertise barrier and accelerates the curation and validation time by orders of magnitude. Moreover, it enables automated interpretability of the neural spiking data, which cannot be achieved by any conventional methods. SpikeAgent presents a paradigm shift in processing signals for neuroscience and brain-computer interfaces, while laying the ground for AI agent-augmented science across various domains.

## Main text

Spike sorting, the process of isolating and classifying single-cell action potentials from extracellular recordings, is a fundamental step in analyzing neural activity^1^. The conventional spike sorting pipeline (**Fig. 1a**) typically involves several key steps: first, preprocessing raw extracellular recordings to filter out noise and isolate relevant frequency bands; second, detecting potential spikes by identifying threshold crossings or similar criteria; third, extracting features (e.g., waveform shape, interspike intervals (ISI), or principal components) from the detected spikes; fourth, clustering these features to assign spikes to individual neurons; and finally, validating the results through post-clustering manual curation^1^. Although this pipeline is widely used in neuroscience, it faces growing challenges related to efficiency, consistency, and scalability as neural datasets rapidly increase in size and complexity^2, 3^.

**Fig. 1:**
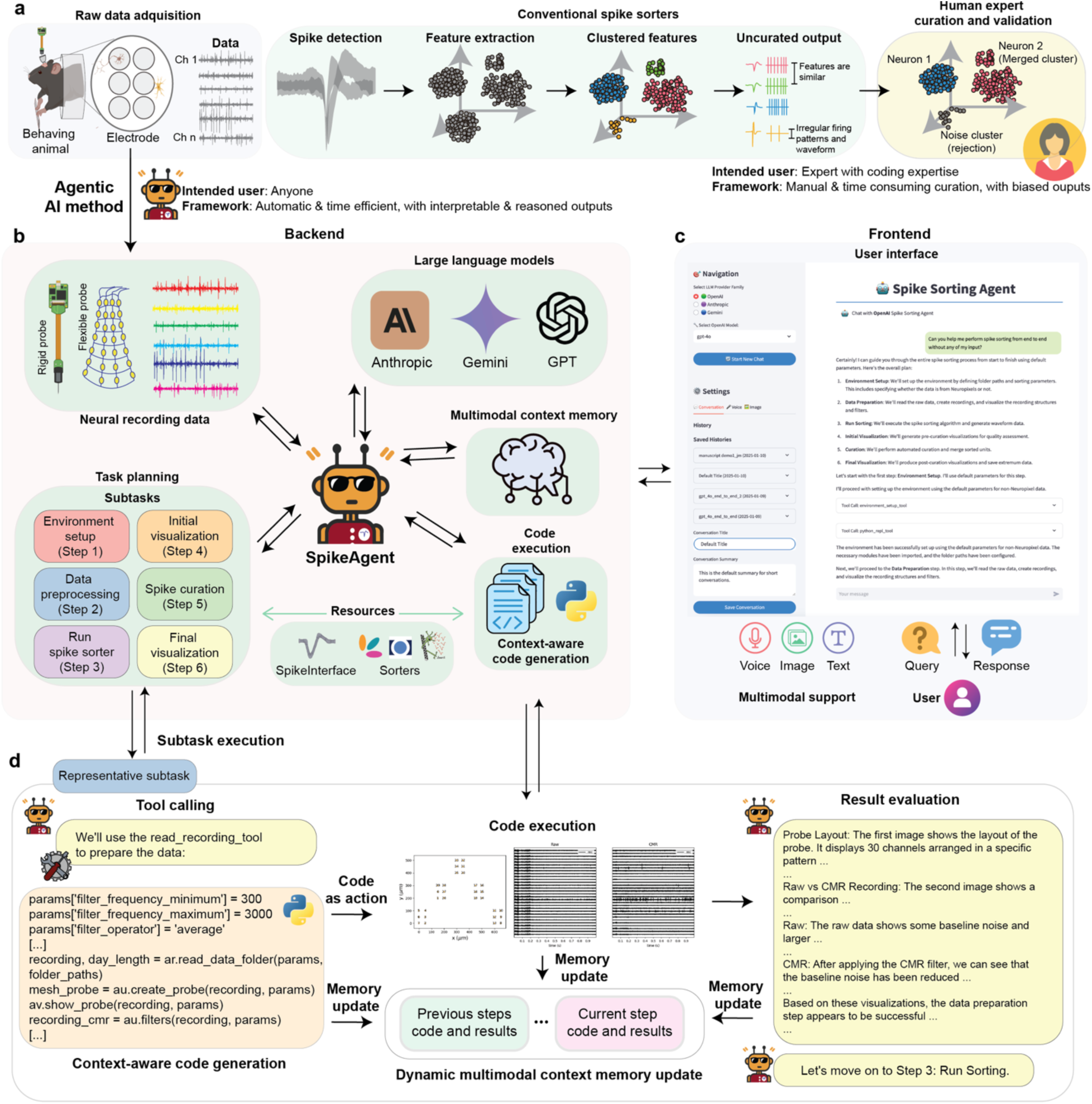
Overview of SpikeAgent framework for spike sorting. (**a**) Schematics showing that conventional spike sorting follows a sequential pipeline, including neural data acquisition, filtering, spike detection, feature extraction, clustering, and manual curation. This process requires expert users with coding expertise and extensive manual efforts, which is highly time-consuming and susceptible to user biases. **(b)** Schematics shows that SpikeAgent introduces an AI agent-driven framework that automates and streamlines the spike sorting process. It integrates neural recording data from various neural recording technologies and employs an AI agent for task planning. The backend incorporates large language models (LLM) (e.g., Anthropic, Gemini, GPT), a multimodal context memory, context-aware code generation, and automated code execution, enabling efficient and interpretable spike sorting. **(c)** Representative example showing that the SpikeAgent user interface supports multimodal interactions, allowing users to input text, voice, and image queries. This intuitive query-response system and history tracking enhance usability and reproducibility. **(d)** Illustration of SpikeAgent’s subtask execution process. Upon receiving a task, the SpikeAgent generates context-aware code for each specific subtask, executes the code, and produces visual outputs. The agent evaluates these results, updates its multimodal context memory with current and past task information, and determines the next step in the workflow. This iterative process of code generation, execution, evaluation, and memory updating enables SpikeAgent to perform complex spike sorting tasks autonomously and efficiently.

The rapid advancement of neuroelectronic technologies, such as high-density Neuropixels for large-scale neural recording^4, 5^ or flexible electrodes for long-term stable recording^6, 7^, have dramatically increased the volume of neural data. While various spike sorting frameworks have been developed^8–11^ to automate spike detection and clustering, manual curation remains essential for ensuring human-level accuracy. This process is labor-intensive, time-consuming, and technically demanding, requiring substantial computational and domain expertise. In addition, the absence of standardized curation guidelines leads to reproducibility issues, making it difficult to achieve consistent results across different studies^12–14^. As neural datasets continue to grow in scale and complexity, traditional spike sorting methods are becoming inadequate for large-scale, high-throughput neural data analysis.

Recent advances in artificial intelligence (AI), particularly large language models (LLMs)^15, 16^ and multimodal AI systems^17^, offer promising solutions to address these challenges^18^. While LLM-based AI agents^19^ has begun to emerge in fields such as chemistry^20, 21^ and material science^22^, their application to neuroscience, especially for spike sorting, remains unexplored. To bridge this gap, we introduce SpikeAgent (**Fig. 1b-d**), an AI agent that leverages multimodal LLM capabilities to fully automate and standardize spike sorting. Our system can operate either fully-autonomously or under human-in-the-loop supervision, generating comprehensive reports that provide detailed reasoning for each curation decision.

SpikeAgent streamlines the entire spike sorting workflow through a conversation-driven interface. Users interact with the agent using natural language, while the agent integrates advanced natural language processing, task planning^23^, context-aware code generation^24^, and visual data interpretation^25^ to handle the task. Unlike traditional methods that rely on rigid, pre-defined workflows, SpikeAgent leverages the reasoning capabilities of LLMs to dynamically interact with data, making context-aware decisions throughout the sorting process. This adaptability allows SpikeAgent to effectively manage the inherent complex conditions of neural recordings, ensuring consistent and high-quality results across diverse datasets.

A key advantage of SpikeAgent is its ability to provide interpretable reasoning through multimodal visual data interpretation, a feature absent in conventional sorting techniques^26^. This interpretability enhances result validation and transparency in the automated process, addressing a critical need for standardized analysis in neuroscience. Moreover, this system enhances reproducibility through centralized memory, aligning with the growing emphasis on transparency and reliability in neuroscience research. By overcoming accessibility barriers and eliminating the need for coding expertise, SpikeAgent democratizes neural data analysis, making it accessible to researchers from diverse backgrounds.

SpikeAgent is designed to accommodate a variety of neural recording technologies, integrates multiple multimodal LLM backends (e.g., GPT^27^, Anthropic^28^, Gemini^29^) along with specialized tools for processing data from flexible electrodes, Neuropixels, and virtually any other recording platform. We systematically evaluate its performance across different LLM families. This comparative analysis validates our approach’s effectiveness and provides insights into the strengths and limitations of various LLM models in the context of long-range task execution and code generation. Benchmarking results demonstrate that SpikeAgent, powered by vision-language models (VLMs), significantly outperforms human experts in both processing speed and curation consistency. By reducing typical curation times from hours to minutes, SpikeAgent transforms previously unmanageable workflows into scalable, efficient processes.

SpikeAgent offers an automated, highly scalable, and interpretable solution to the longstanding challenges of spike sorting. By significantly reducing time and expertise requirements and eliminating coding prerequisites, SpikeAgent accelerates neural data analysis and enables new insights into brain function. As recording hardware continues to advance, capturing ever-larger neural populations with increasing precision and scale, it will soon become impractical to rely solely on human curation. SpikeAgent provides the foundation for a new era of AI agent-driven, automated, standardized, and interpretable neural data analysis, enabling neuroscience to fully leverage the potential of modern electrophysiological recording technologies.

## Results

### SpikeAgent system architecture

SpikeAgent integrates various components into a cohesive system that fully automates the entire spike sorting process. **Figure 1** illustrates the architecture of this system, which consists of two primary components: the backend and the frontend user interface (**Fig. 1b-c**).

The backend (**Fig. 1b**) serves as the core processing unit of SpikeAgent, interfacing with multiple multimodal LLM providers (e.g., GPT^27^, Anthropic^28^, Gemini^29^), which act as the system’s reasoning core and planning engine. These LLMs operate alongside a multimodal context memory, allowing for real-time, dynamic updates and retention of multimodal information throughout the spike sorting process. This multimodal context memory is crucial for an AI agent to make context-aware decisions at each step of the spike sorting process. SpikeAgent processes various input modalities (voice, image, text) and manages the flow of information between different system components. A key feature of the LLMs is their ability to perform task planning and orchestration, allowing SpikeAgent to break down the spike sorting process into manageable subtasks, including environment setup, data preprocessing, spike sorter running, initial visualization, spike curation, and final visualization. This modular approach ensures a systematic, comprehensive, and scalable analysis of neural recording data. Another critical feature is context-aware code generation, which enables SpikeAgent to dynamically generate and execute code tailored to each dataset and experimental setup. This is enabled by the interaction between the SpikeAgent, the multimodal context memory, and the code execution module.

The frontend (**Fig. 1c**) provides an intuitive, chat-based interface, allowing researchers to interact with SpikeAgent using text, image, and voice inputs (**Extended Data Fig. 1**). This design makes the system accessible to researchers with varying levels of technical expertise. The frontend incorporates features such as model provider selection, multimodal input options, and conversation history management. Users can easily switch between different LLM backends, input data through text, voice, or images, and save or load previous conversations. This comprehensive interface not only facilitates ease of use but also enhances reproducibility and reusability of results.

The system leverages context-aware Python code to consolidate all actions into a unified action space. During each subtask execution (**Fig. 1d**), SpikeAgent automatically generates code, which is then executed in a Python environment. For instance, during data preparation, the SpikeAgent automatically selects the appropriate tool (e.g., read_recording_tool) and generates code with optimized parameters for data filtering and preprocessing. This execution produces visual outputs crucial for downstream analysis. The dynamic multimodal context memory, a key feature of the SpikeAgent, continuously updated throughout the process, retains information from previous steps and incorporates newly generated results, enabling a coherent and contextually aware workflow. After completing each subtask, the agent evaluates the outcomes, analyzing both visual and textual elements, such as probe layout, raw vs. common mode reference (CMR) recordings, and the effectiveness of noise reduction techniques, ensuring that all steps contributed to an optimized spike sorting process. This capability allows SpikeAgent’s to interpret complex neural recording data and make informed decisions for subsequent steps, including self-debugging and autonomous adjustment of its approach when necessary.

Upon completing all subtasks, SpikeAgent generates a comprehensive final output, detailing the curation results of the spike sorting process. The system’s architecture leverages the strengths of multimodal LLMs while maintaining a user-friendly interface, thereby increasing the efficiency and accessibility of spike sorting in neuroscience research. This integrated approach allows SpikeAgent to handle the complexities of spike sorting with minimal human intervention, transforming raw neural recording data to sorted spike data with high reproducibility and standardization.

### Fully automatic end-to-end and human-AI collaboration for spike sorting

SpikeAgent addresses the limitations of traditional spike sorting methods, which rely heavily on human intervention, by offering a versatile, AI-driven solution for users regardless of their coding expertise. It provides two primary execution modes: a fully automated, end-to-end workflow that requires no human input (**Fig. 2a**) and a human-in-the-loop mode that allows for interactive, user-guided analyses (**Fig. 2b**). This dual functionality enables researchers to choose between a streamlined, hands-off automation approach or a customized, user-guided workflow, depending on their specific needs (i.e. specific recording hardware).

**Fig. 2:**
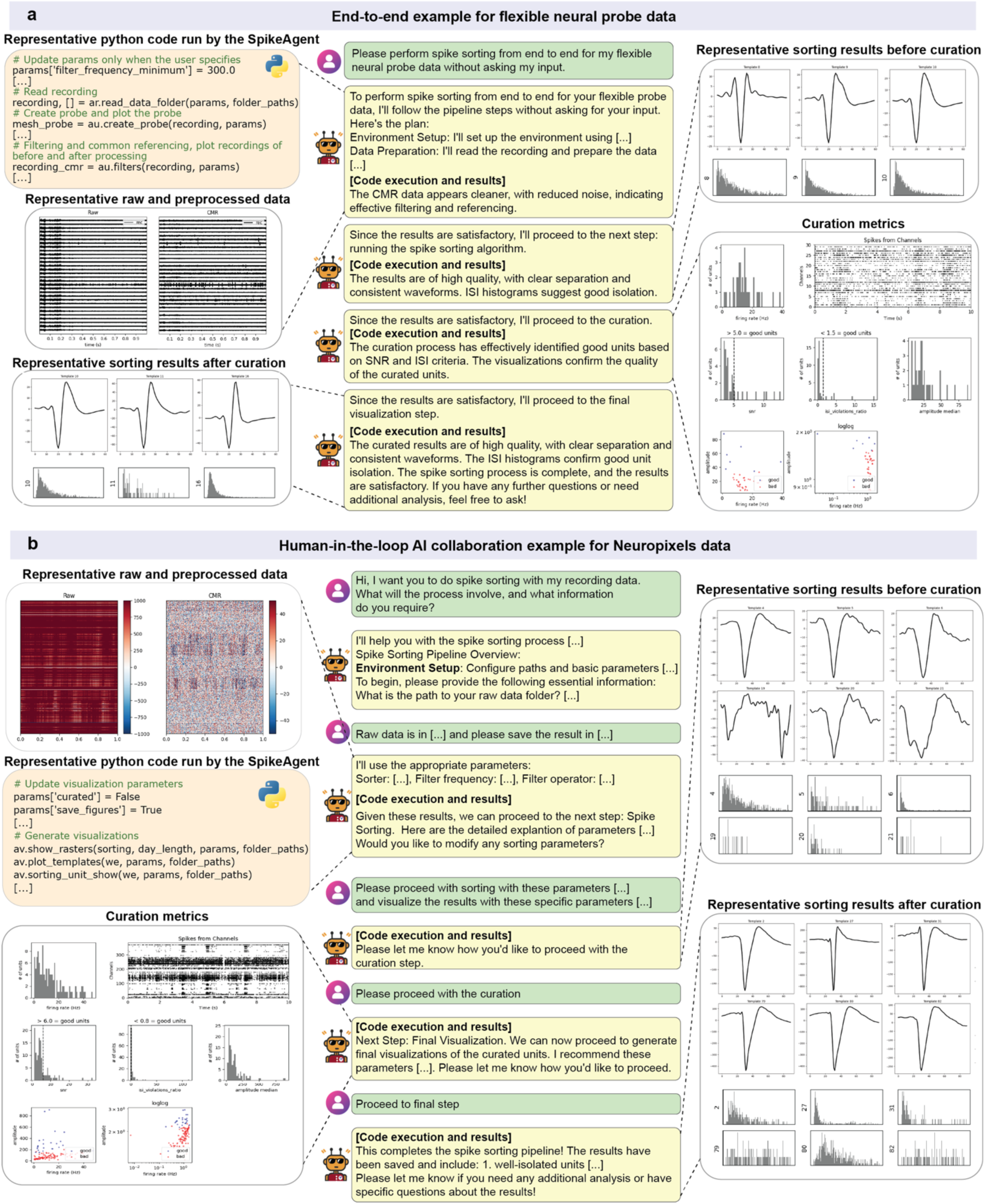
End-to-end and human-AI agent collaboration in SpikeAgent. **(a)** Autonomous spike sorting for flexible neural probe data. SpikeAgent executes the entire spike sorting pipeline without user intervention. The workflow begins with automated task planning for the entire spike sorting pipeline, followed by Python code execution (top left), where parameters are configured, and raw neural data is preprocessed. Representative raw and preprocessed data highlight the result of preprocessing, including significant baseline noise reduction. Zoomed-in views (top right) illustrate individual spikes before curation, emphasizing the effectiveness of the sorting pipeline. After curation, results show enhanced isolation of single units, validated by inter-spike interval (ISI) histograms and well-defined waveforms (bottom left). Key curation metrics, including false-positive rates and cluster quality, confirm the robustness of SpikeAgent’s spike sorting (bottom right). **(b)** Human-AI agent collaboration for Neuropixels data. This workflow begins with user input specifying data requirements and preferences, followed by SpikeAgent’s execution of spike sorting in a guided, stepwise manner. Preprocessed data (top left) shows both raw recordings and noise-reduced outputs, with zoomed-in sections highlighting noise removal and enhanced spike visibility. Sorting parameters are iteratively adjusted through user feedback, producing sorting results before (top right) and after (bottom right) curation. Results illustrate enhanced unit isolation, validated through ISI histograms and waveform separation, confirming successful noise reduction and cluster refinement. Curation metrics (bottom left), provide a transparent evaluation of the final curated results, demonstrating the agent’s ability to collaborate with users effectively while ensuring high-quality outcomes.

In the fully automated mode, SpikeAgent executes the entire spike sorting workflow using simple natural language input **(Fig. 2a)**. The process begins with the user providing a high-level natural language request along with neural recording data (**Methods**). For example, when asked, “*please perform spike sorting from end-to-end for my flexible electrode recording data without asking my input.*”, SpikeAgent automatically initiates its task-planning process (**Fig. 2a**), breaking down the spike sorting workflow into a series of subtasks, including environment setup, data preprocessing, spike sorting, curation, and final visualization. This ability to autonomously interpret and translate complex requests into structured workflows shows the SpikeAgent’s intelligence and adaptability. Once the workflow is initiated, SpikeAgent automatically executes each subtask sequentially, generating and running Python code for each step. During data preparation, for instance, the agent reads the recording, displays the geometry layout of the neural probe, and applies filtering and common referencing to optimize the signal quality (**Methods**). After each subtask, the agent evaluates the intermediate results and automatically proceeds to the next step, while providing updates to the user on its progress. Throughout the process, SpikeAgent generates visual outputs such as representative raw and preprocessed data, sorting results before and after curation, and key curation metrics. These outputs include waveform plots, ISI histograms, and various quality metrics, which provide a thorough assessment of the spike sorting pipeline. The final output demonstrates high-quality sorting results, with clear spike separation and consistent waveforms, confirming the effectiveness of the fully automated process (**Fig. 2a**).

For users who prefer a more interactive approach, SpikeAgent also supports a human-in-the-loop collaboration mode (**Fig. 2b and Extended Data Fig. 2**), as exemplified in a use case with Neuropixels data (**Methods**). This mode enables users to provide iterative feedback and specify custom requirements during the spike sorting tasks. The workflow begins with the user inquiring about the spike sorting procedure, prompting SpikeAgent to outline the steps and request relevant information. As the workflow progresses, the user can adjust parameters, request specific visualizations, and make decisions at key points, such as proceeding with curation or final visualization steps. For example, after initial data preprocessing, the agent presents intermediate results and asks the user if further adjustments of any sorting parameters are needed. The user can then provide specific instructions, such as “*Please proceed with sorting with these parameters […] and visualize the results with these specific parameters […]*”. This iterative exchange continues through the curation and final visualization steps, with SpikeAgent dynamically executing code and generating results based on user feedback at each stage. By offering detailed explanations of parameters and results throughout the analysis, this mode enhances transparency and user control, ensuring that analysis aligns with specific research requirements.

Beyond its automation and interactive capability, SpikeAgent’s versatility extends to various interaction patterns (**Extended Data Fig. 3**). For example, it can provide knowledge-based responses (**Extended Data Fig. 3a**), answering user queries about spike sorting concepts to help researchers better understand the process. In addition, SpikeAgent can perform computation-augmented responses (**Extended Data Fig. 3b**) for specific calculations and generate visualization-driven responses (**Extended Data Fig. 3c**) to create complex, task-specific plots on demand. This versatility allows SpikeAgent to adapt to different data types, user expertise levels, and analysis requirements, making it a powerful tool for spike sorting and neural data analysis across various scenarios.

### Comparative analysis of different LLM backends

We evaluated the performance and capabilities of SpikeAgent across different multimodal LLMs, conducting a comparative analysis using GPT, Anthropic, and Gemini as the underlying backend models (**Fig. 3a**). Each instance of SpikeAgent with different backend LLM (**Methods**) and prompt instructions, received the same user input: “*Hello. perform the entire spike sorting pipeline of non_neuropixel type, without asking me for details of the parameters. Use the default for all steps*.” (**Methods**). This approach allows us to systematically assess the relative strengths and weaknesses of each LLM in task planning, instruction following, and code generation for different key steps in spike sorting. By standardizing the input and task requirements, we aimed for a fair comparison across all tested models.

**Fig. 3:**
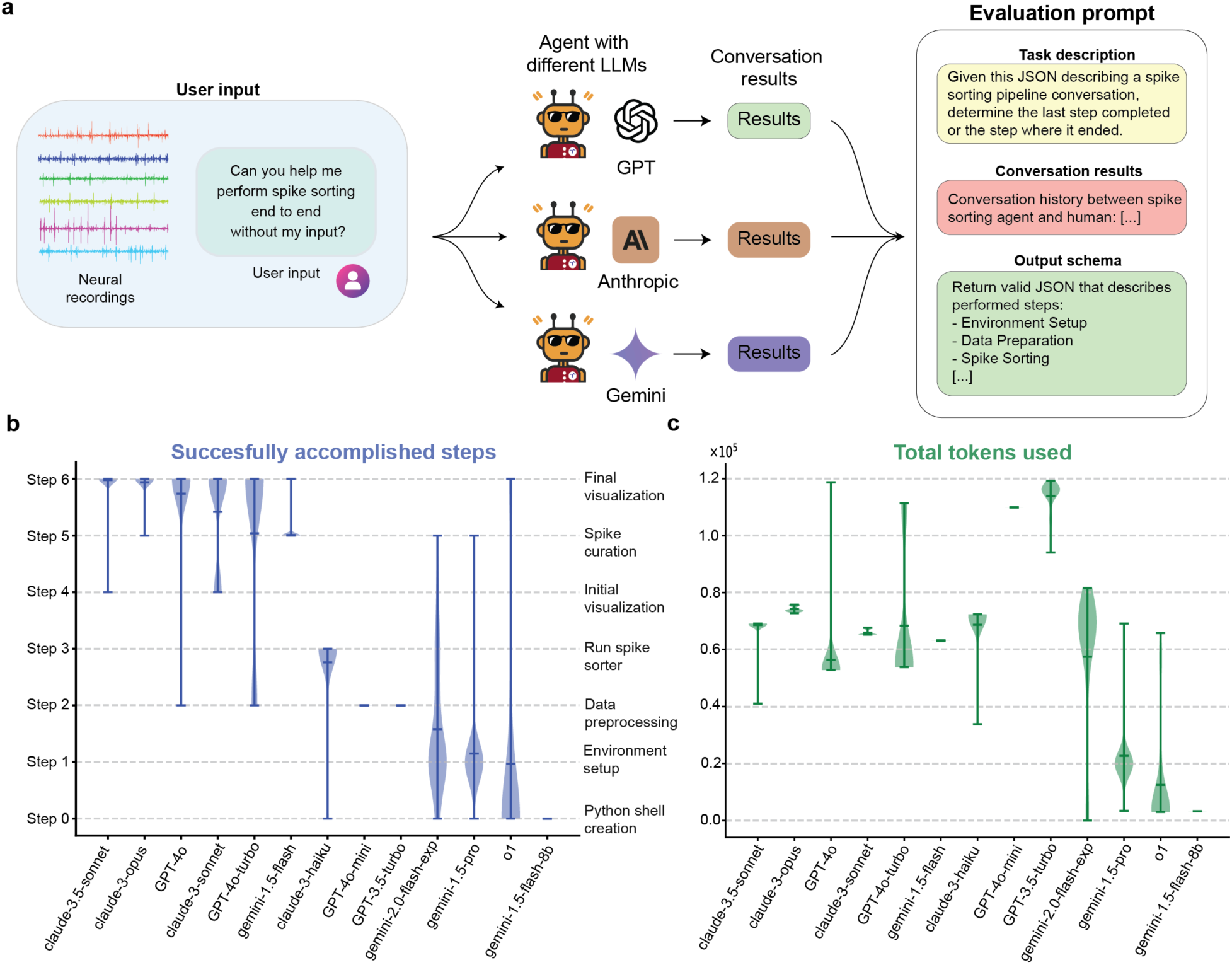
Performance evaluation of SpikeAgent’s LLM backends for autonomous spike sorting. (**a**) Evaluation workflow comparing different LLM backends (GPT, Anthropic, and Gemini) based on their ability to execute spike sorting tasks through the combination of instruction following, task planning, and code generation. The same neural recording data and user input are processed by SpikeAgent, which generates corresponding code. The performance of each LLM backend is evaluated using task-specific prompts that consider detailed task descriptions, conversation history, and output schemas. (**b**) Benchmark of successfully accomplished steps without code failure, comparing the reliability of each LLM backend in executing tasks. (**c**) Total tokens used by each model, measuring computational resource consumption and efficiency. All violin plots represent the distribution of each data over different seeds (N=100), with width indicating frequency. Horizontal lines show the mean, maximum, and minimum.

To analyze the results, we employed an LLM-as-a-judge approach^30^, utilizing an evaluation framework centered around a specific prompt (**Fig. 3a**, right panel). This evaluation method involves analyzing the conversation results between the human user and different LLM-powered SpikeAgents to determine the last successfully completed step or the point of failure in the spike sorting pipeline. By tracking where each model failed, we could evaluate how far each model progressed through the spike sorting pipeline and identify any coding points of failure or divergence. This approach not only quantified task completion but also provided insights into the stability and consistency of the models across repeated trials.

The completion rate analysis (**Fig. 3b**) reveals substantial differences in LLM performance to successfully complete seven (Step 0-6) sequential steps in the spike sorting pipeline (**Fig. 1b**). Among the tested models, Claude-3.5-sonnet, Claude-3-opus, GPT-4o, and Claude-3-sonnet emerged as the top performers, consistently completing the entire pipeline with high stability across trials. These models achieved robust performance, demonstrating their capability in handling the pipeline’s complexity. In contrast, other models, such as Gemini-1.5-flash-8b and o1, exhibited lower task completion rates and higher variability, limiting their reliability for spike sorting. For instance, Gemini-1.5-flash-8b often failed to even create the Python shell required to execute any of the pipeline tasks, showing inability to progress beyond the initial stages. Consequently, for subsequent analyses and practical usage, we focused exclusively on the top-performing models, particularly those equipped to handle multimodal (visual) inputs, which is a critical requirement for spike sorting workflows.

Beyond task completion, we also analyzed token usage across different LLMs (**Fig. 3c**), revealing substantial differences in their computational cost. Each model family employs a unique approach to token usage computation, reflecting their different trade-offs between verbosity, efficiency, and performance. Among the top-performing models, Claude-3-opus exhibited the highest token usage, followed closely by Claude-3.5-sonnet. While these models demonstrated high task completion rates, their elevated token consumption suggests their reliance on verbose responses to achieve high performance. Conversely, GPT-4o achieved competitive task completion rates with lower token usage, demonstrating their efficiency. In contrast, Gemini models, such as Gemini-1.5-flash-8b and o1, exhibited minimal token usage but consistently failed to complete the spike sorting workflow. The differences in token consumption reflect distinct architectural trade-offs across model families. These differences also stem from their distinct methods of token accounting, with some prioritizing contextual verbosity while others opt for minimalistic responses. This highlights the trade-off between maximizing task completion and ensuring computational efficiency, a critical consideration for scalability and deployment in large-scale applications.

### Automatic explainable spike curation enabled by VLM-powered SpikeAgent

One key strength of LLMs is their reasoning capability. Traditional spike sorting curation often relies on subjective criteria and manual curation, leading to inconsistencies across studies and among human experts. This lack of standardization limits the robustness and transparency of neural recording data analysis. In addition, manual curation rarely provides detailed reasoning for accepting or rejecting individual spikes, making it difficult to validate decisions or identify potential biases in the curation process. These issues become increasingly problematic as modern neural recording techniques generate larger and more complex datasets.

To address these limitations, we explored whether SpikeAgent could address the lack of standardization and interpretability in traditional spike sorting by leveraging the advanced visual and reasoning capabilities of VLMs^25^ to enable automated and explainable spike curation (**Fig. 4a**). By integrating both textual and visual inputs, the VLM applies detailed curation criteria, scoring guidelines, and examples of “good” and “noisy” spikes to inform its decisions. SpikeAgent uses in-context few-shot learning, providing the VLM with a rich context for each classification task. This context includes a comprehensive task description, specific curation criteria, quantitative scoring guidelines, and visual examples of both good and noisy spikes. This approach allows the VLM to make informed decisions by comparing input query images against provided examples and criteria, enabling quick adaptation to new datasets and curation standards without model retraining.

**Fig. 4:**
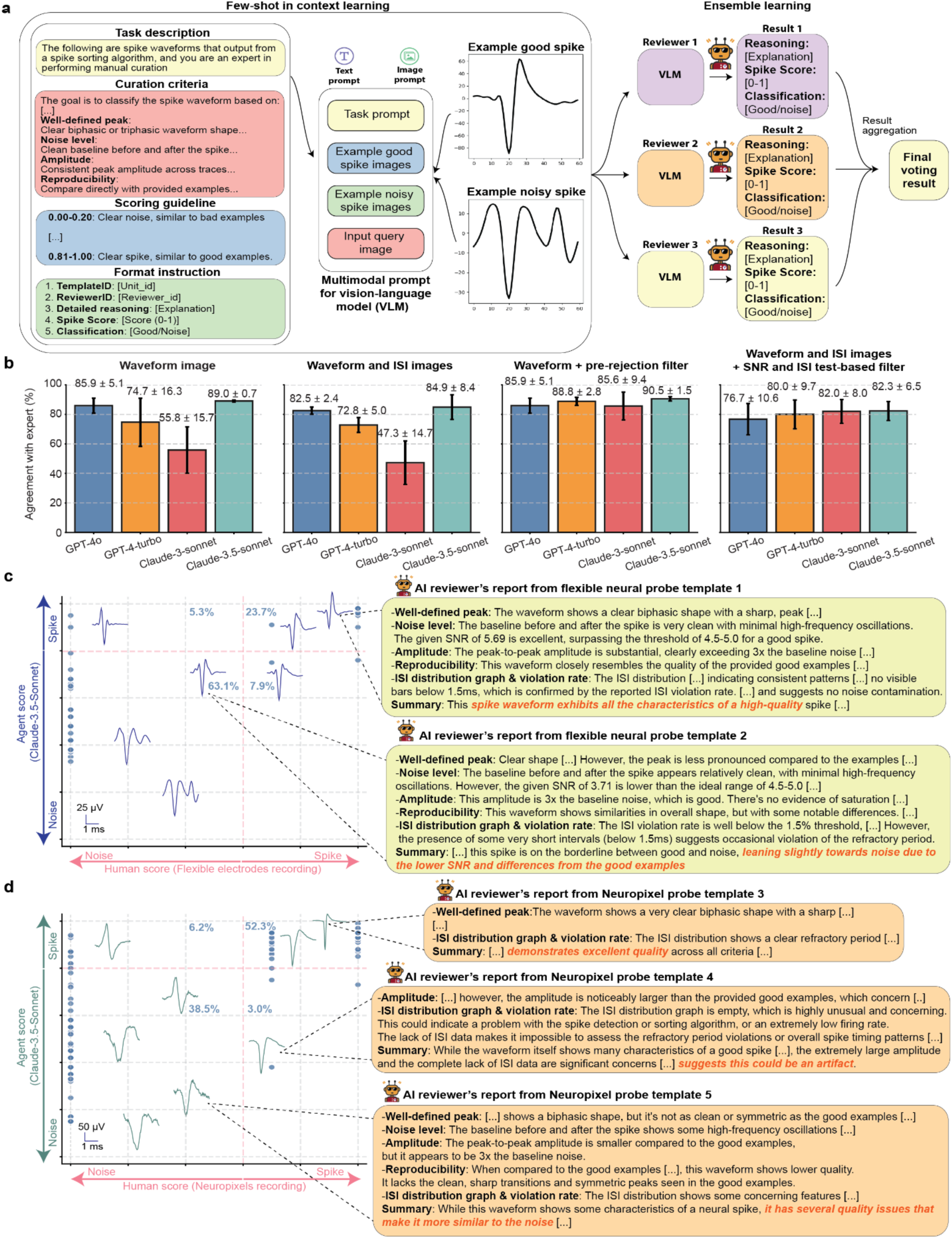
Few-shot in-context learning framework using vision-language model (VLM) for automated explainable spike curation. **(a)** Few-shot in-context learning framework for spike waveform curation using a VLM. The task is guided by predefined curation criteria and scoring instructions. Multimodal prompts include waveform images, textual descriptions, and examples of “good” and “noisy” spikes as references. The VLM evaluates each waveform based on criteria such as peak shape, noise level, amplitude consistency, and ISI metrics. Multiple independent VLM reviewers classify waveforms as “Good” or “Noise”, providing explanations for their decisions. The final classifications are determined through ensemble learning, aggregating outputs from multiple reviewers. **(b)** Accuracy comparison of different VLM input modalities for spike curation benchmarked with human expert’s curation consensus. Bar plots show model performance under four conditions: waveform images alone, waveform with ISI images, waveform with ISI images after pre-rejection filtering, and waveform images with ISI images supplemented with text-based ISI and SNR metrics. Data are mean ± s.d. **(c-d)** VLM-based spike curation for neural recordings from flexible electrode (**c**) and Neuropixels (**d**). Scatter plots show alignment between VLM predictions and human-curated scores. Representative waveform examples illustrate VLM’s reasoning in evaluating spike features, explaining how it determines waveform quality.

By utilizing chain-of-thought reasoning^31^ and ensemble decision-making^32^, VLM-powered SpikeAgents provide interpretable and transparent justifications for their outputs, enabling reliable and reproducible curation. The ensemble approach employs multiple independent VLM “reviewers”, each of which analyzes individual spikes separately (**Fig. 4a**). Each reviewer provides its detailed reasoning, spike scores, and classification, which are then aggregated to produce a final voting result with enhanced robustness and reliability for classification. This multi-reviewer system mimics the collaboration between multiple human experts for manual curation, while leveraging the consistency and scalability of AI. This combination of few-shot learning, VLM-enabled multimodal input processing, and ensemble decision-making allows SpikeAgent to classify spikes, identify subtle patterns, and adapt to a wide range of datasets. This approach transforms the spike sorting workflow into a scalable, automated, and highly interpretable process, establishing a new standard for neural data analysis.

To systematically evaluate the impact of different input modalities on agreement with human experts consensus and interpretability, we evaluated four multimodal strategies: (1) waveform images only, (2) waveform images with ISI distribution images, (3) waveform images with pre-rejection filtering, where predefined code functions removed spikes with low SNR or high ISI violation rates, and (4) waveform images with ISI images and text-based SNR and ISI values, which act as filtering following guidelines described in the prompt (**Methods**). These strategies enabled a systematic exploration of how different input combinations and ensemble reasoning contribute to spike classification and curation accuracy and interpretability.

We conducted benchmarking using two neural recording datasets: flexible neural probe data and Neuropixels data (**Methods**). The agreement with human experts’ comparison (**Fig. 4b**) reveals that waveform images with pre-rejection filtering achieved the highest agreement, consistently exceeding 85% agreement with human experts’ consensus. This pre-rejection step reduces the impact of erroneous inputs, improving classification consistency across different VLMs. Simpler approaches, such as waveform images alone, also performed well, with Claude-3.5-sonnet and GPT-4o achieving 89.0 ± 0.7% and 85.9 ± 5.1% agreement with human expert consensus, respectively. However, adding ISI images without pre-defined filtering sometimes reduces accuracy, suggesting that this modality may introduce additional complexity that not all models can handle efficiently. These findings emphasize that filtering, whether pre-applied through code or incorporated via task prompts, generally improves classification accuracy across models while retaining flexibility to perform well in simpler configurations. We observed that when using code-based filtering, the reasoning becomes less informative, since the SpikeAgent has less information in its context to use. However, in the fourth modality when using the SNR and ISI metrics as filtering via text inputs, the reasoning became more informative.

To emphasize these findings, we studied the reasoning reports generated by SpikeAgent and compared them with the human experts in the flexible neural probe dataset (**Fig. 4c**). For example, template 1, a true spike, was classified as a spike due to its distinct biphasic waveform, low noise level, and amplitude consistent with good spike examples. The addition of ISI images and SNR and ISI text filters improved the model’s ability to integrate temporal and statistical information, supporting the reasonings aligned with human experts. For example, on template 1, one of the reviewer’s reports suggested “*the baseline before and after the spike is very clean […] The given SNR of 5.69 is excellent*” or “*The ISI distribution […] indicates consistent patterns, […] which is confirmed by the reported ISI violation rate*”. This also happened in template 2, classified as noise. This template was identified based on low SNR and deviations from the ideal spike characteristics. One of the AI agent reviewers justified its classification by “*Clear shape […]. However, the peak is less pronounced compared to the examples*” and “*the SNR of 3.71 is lower than the ideal range of 4.5-5.0*”. These findings demonstrate that SpikeAgent can provide an additional interpretability layer beyond conventional spike sorting methods. Overall, this VLM modality achieved a very high agreement with human experts of 86.8% (63.1% true noises, 23.7% true spikes) as well as providing an additional unique layer of reasoning that is unapproachable by any of the other modalities, at the cost of minimal extra computations.

On the other hand, for Neuropixels probe data, we evaluated VLM performance using waveform images combined with ISI images, without additional filtering (**Fig. 4d**). This configuration also showed strong agreement with human curation. Template 3, a true spike, was classified based on its clear biphasic waveform, low noise level, and high reproducibility across traces. In contrast, template 5, a true noise, was correctly rejected due to ambiguous peaks and significant noise, characteristics consistent with noise examples. The VLM achieved an overall agreement with the human expert’s consensus of 90.8%, with 52.3% true spikes and 38.5% true noises. One misclassified example was template 4. The VLM classified it as noise, while the human expert identified it as a spike. The misclassification occurred because the AI lacked sufficient ISI information and found discrepancies with its few-shot training examples. In cases of uncertainty, the VLM favored caution by classifying ambiguous signals as noise. This configuration demonstrates that waveform and ISI images serve as complementary inputs, enabling accurate classification without requiring pre-rejection filtering, which optimizes the efficiency, since no further computation is required to make any decisions.

The SpikeAgent performed consistently across different high-density electrophysiological recordings, achieving high agreement with human experts across both flexible neural probe and Neuropixels datasets (**Table 1 and Table 2**). To further evaluate the VLMs adaptability and demonstrate the impact of providing few-shot examples, we tested zero-shot in-context learning, where models received identical task guidelines but no labeled examples for reference (**Extended Data Fig. 4**). The zero-shot approach showed substantial variation across models. For example, GPT-4o achieved the highest agreement with human experts using waveform images alone, while GPT-4-turbo and Claude-3.5-sonnet performed worse. Adding ISI images had mixed effects: GPT-4-turbo improved but GPT-4o dropped, suggesting that, like the few-shot scenario, ISI images introduce complexity that not all models handle well (**Extended Data Fig. 4**).

**Table 1:** Table showing the agreement between each VLM model and human expert consensus in spike curation for the flexible neural probe dataset.

| **Flexible neural probe** | GPT-4o | GPT-4-turbo | Claude-3-Sonnet | Claude-3.5-Sonnet |
| --- | --- | --- | --- | --- |
| Waveform images | 89.5 | 64.2 | 44.7 | 89.5 |
| Waveform and ISI images | 84.2 | 76.3 | 38.6 | 78.9 |
| Waveform + pre-rejection filter | 89.5 | 86.8 | 78.9 | 89.5 |
| Waveform and ISI images + SNR and ISI filter (text-based) | 84.2 | 86.8 | 76.3 | 86.8 |

**Table 2:** Table showing the agreement between each VLM model and human expert consensus in spike curation for the Neuropixels dataset.

| **Neuropixels** | GPT-4o | GPT-4-turbo | Claude-3-Sonnet | Claude-3.5-Sonnet |
| --- | --- | --- | --- | --- |
| Waveform images | 82.3 | 86.2 | 66.9 | 88.5 |
| Waveform and ISI images | 80.8 | 69.2 | 57.7 | 90.8 |
| Waveform + pre-rejection filter | 82.3 | 90.8 | 92.3 | 91.5 |
| Waveform and ISI images + SNR and ISI filter (text-based) | 69.2 | 73.1 | 87.7 | 77.7 |

Overall, the zero-shot curation results show lower agreement with human experts, highlighting more the inherent limitations across VLM. These findings demonstrate that while zero-shot vision-based spike curation is feasible, providing in-context examples and structured numerical inputs (e.g., ISI and SNR values) remain essential for robust agreement with human expert and interpretability, particularly in complex neural datasets where these metrics provide crucial context for spike classification.

### SpikeAgent surpasses human curation in speed, consistency, and scalability

Manual spike curation has become increasingly unreliable for modern neuroscience studies due to its slow, inconsistent, and inherently biased nature. The rapid development of high-density, large-scale neural electrode recording technologies has generated datasets that far exceed human curation capabilities in both volume and complexity. Additionally, inconsistencies between human expert curators introduce variability in results across different studies or laboratories, undermining reproducibility and comparability in neuroscience research. To address these limitations, we investigated whether SpikeAgent, leveraging the advanced capabilities of VLMs, could provide a scalable, efficient, and standardized solution.

To evaluate SpikeAgent’s effectiveness, we compared its efficiency and consistency against both human expert and novice human reviewers. **Figure 5** shows three distinct approaches to spike curation: standard VLM curation (**Fig. 5a**), asynchronous VLM curation (VLMa) (**Fig. 5b**), and human curation (expert and novice) (**Fig. 5c**). In standard VLM curation (**Fig. 5a**), multiple VLM reviewers independently process spike curation sequentially, limiting processing speed. In contrast, the VLMa (**Fig. 5b**) curation allows multiple reviewers to process spike curation simultaneously in parallel threads, significantly improving efficiency. Human curation (**Fig. 5c**) includes expert and novice reviewers, grading spikes on a scale of 0 to 1 across four categories: amplitude, noise level, spike shape, and reproducibility.

**Fig. 5:**
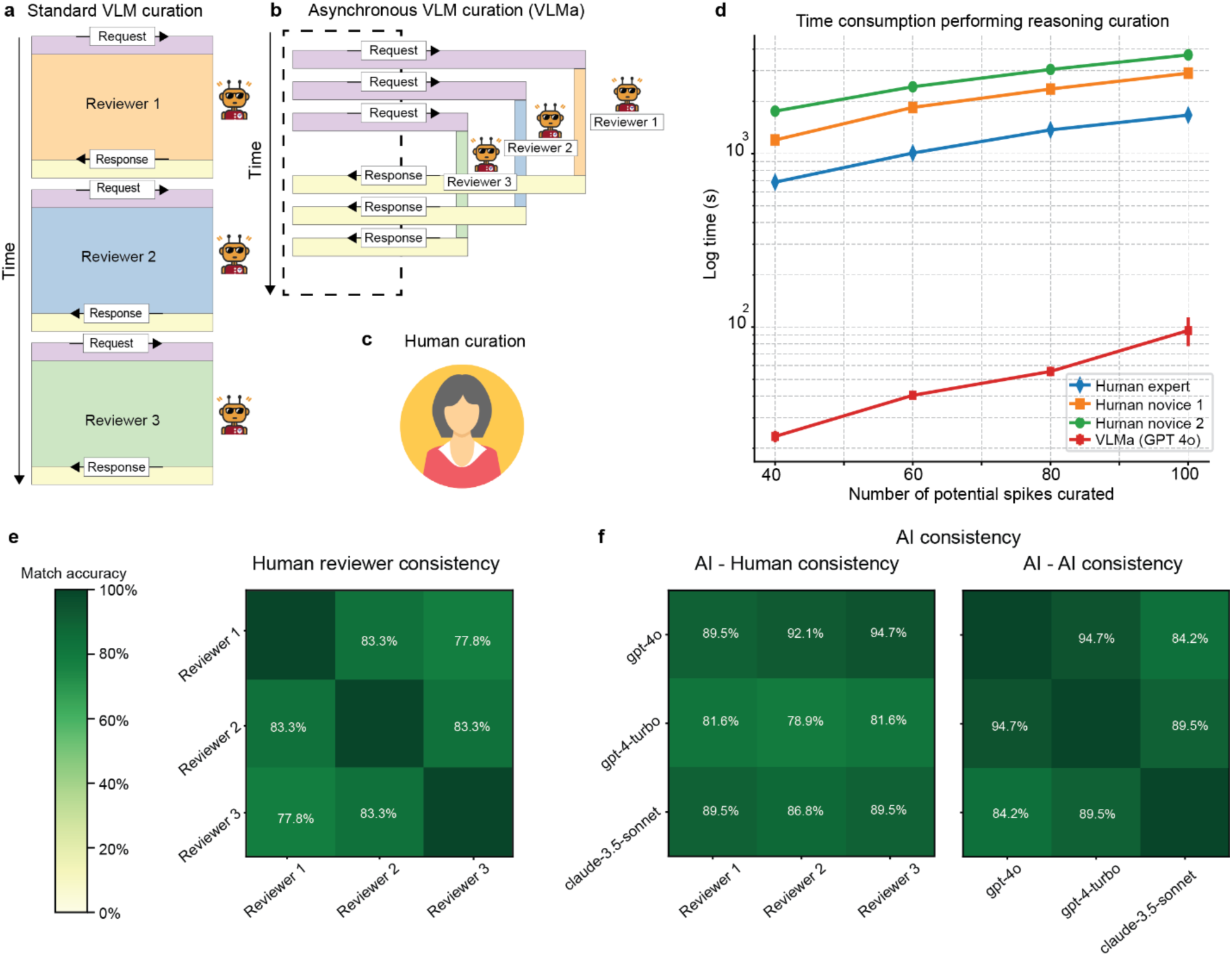
VLM curation efficiency and consistency. **(a)** Standard VLM curation process, where requests and responses are sequentially managed by individual reviewers. (**b)** Asynchronous VLM (VLMa) curation process, allowing parallel input and output from multiple AI reviewers to improve efficiency. (**c**) Human curation processes, including expert and novice reviewers, based on key spike quality parameters. (**d)** Time required for reasoned manual curation across three instances of the VLM asynchronous modality, using GPT-4.0, compared against three human reviewers (one expert and two novices). The dataset used for this benchmark was the Neuropixels. **(e**) Human reviewer (experts only) consistency matrix, showing agreement between individual human reviewers. (**f**) Consistency matrices for AI-human (expert only) comparisons and AI-AI comparisons across multiple models.

Time-efficiency analysis revealed striking disparities (**Fig. 5d**). Human curation, whether by expert or novice, exhibited an exponential time increase as spike counts grew, particularly among novices, due to cognitive and manual constraints. In contrast, VLMa reduced processing time from several hours to just two minutes by leveraging parallel processing potential. Notably, in this study, we conservatively limited batch processing to 40 samples per batch (**Methods**) as a proof-of-concept; therefore, even greater acceleration could be achieved with more OpenAI API keys. This acceleration highlights the advantage of asynchronous parallelization, enabling a scalable approach that could dramatically improve efficiency without compromising interpretability.

Critically, these efficiency gains of VLMa did not come at the cost of classification quality. In fact, human reviewers (in this case, all of them were experts) only showed moderate inter-rater reliability (**Fig. 5e**), with match accuracies ranging from 77.8% to 83.3%, highlighting the inherent subjectivity of manual curation. AI-human consistency (**Fig. 5f**) demonstrated higher agreement, particularly with GPT-4o, which achieved 89.5% to 94.7% alignment with human reviewers. Notably, the AI-AI consistency (**Fig. 5f**) among these models reached up to 94.7%, surpassing the observed range among human reviewers. These findings demonstrate that SpikeAgent’s VLMa configuration eliminates the bottlenecks of manual curation while providing a level of standardization superior to expert-level manual approaches. The framework’s scalable architecture directly addresses the critical needs for reproducibility in large-scale electrophysiological studies.

## Discussion

SpikeAgent introduces a novel LLM-based AI system that automates and standardizes the spike-sorting pipeline across diverse neural recording technologies and algorithms, addressing longstanding challenges in neuroscience research. By integrating multiple multimodal LLM backends with a dynamic multimodal context memory, SpikeAgent substantially improves neural data analysis by enhancing reproducibility and establishing a standardized framework for processing neural recordings across different technologies and methodologies.

SpikeAgent also represents a paradigm shift in how neural recording data can be processed, transforming it into interpretable, actionable signals at a scale unattainable by human curators. This system makes complex neural activity comprehensible and accessible to the broader neuroscience community, ultimately driving new opportunities in systems neuroscience. Specifically, by enabling end-to-end efficient spike-sorting workflows driven by simple natural language inputs, SpikeAgent democratizes access to advanced neuroscience techniques, empowering researchers regardless of their coding expertise or specialized knowledge of spike-sorting algorithms. In large-scale datasets containing millions of spikes or when comparing multiple datasets across different technologies and platforms, human curation becomes impractical and unreliable. SpikeAgent overcomes these challenges by offering a scalable, unbiased, and standardized solution, ensuring consistency and efficiency in spike sorting analysis.

Our findings demonstrate that SpikeAgent’s VLM-driven spike curation consistently achieves high accuracy. Benchmarking against human expert curation across diverse recording types and sorting algorithms revealed a high level of agreement between VLM outputs and human classifications, with observed discrepancies typically within 10-20%, a range that aligns with inter-human variability in such tasks^12^. Additionally, VLM-driven scoring system offers a continuous classification scale (0–1), enabling flexibility curation strategies. For instance, spikes with mid-range scores (e.g., 0.5–0.7), often categorized as noise, can be revisited for closer inspection as they may represent ambiguous signals. Similarly, borderline high scores (e.g., 0.81–0.85) can be re-evaluated based on user-defined thresholds. This adaptability ensures that advanced users can fine-tune the curation process based on specific experimental requirement or specialized recording technologies, enhancing both reliability and usability of the system.

SpikeAgent’s efficiency is particularly noteworthy, with VLMa curation significantly outperforming both standard VLM and human curation in terms of time efficiency, achieving substantial acceleration through parallel processing. This efficiency again enables scalable reasoning without compromising interpretability or accuracy, potentially transforming the pace of large-scale neural data analysis in neuroscience research.

Despite its strengths, several limitations remain for the AI agent field to further explore. First, we observed that SpikeAgent’s curation decisions are sensitive to factors such as prompt design (e.g., instructing the model to act as a “strict” vs. a “generous” reviewer), input modality (e.g., waveform images alone vs. waveform + ISI images), and prompting strategy (zero-shot vs. few-shot). For instance, providing well-structured examples (e.g., labeled waveforms) in the prompt significantly improved stability and interpretability compared to purely zero-shot setups. Second, while we tested SpikeAgent on flexible neural probes and Neuropixels probes, its performance on a broader range of electrode arrays and across different species remains to be explored. Finally, certain LLMs (e.g., GPT-o1) struggled with function-calling and managing complex workflows, indicating that not all LLMs are equally effective for code-based tasks. These challenges highlight the need for prompt engineering to account for different input modalities, comprehensive benchmarking across diverse neural recording technologies, and carefully LLM selection to ensure robust handling of complex steps and errors.

Looking ahead, the development of SpikeAgent opens exciting opportunities for future research and applications. For example, when applied to new neural recording technologies lacking any spike-labeled data, SpikeAgent could still offer valuable insights by combining zero-shot classification with established analytical frameworks. An adaptive approach could involve initial noise filtering using established quality metrics, paired with probabilistic confidence scores to estimate spike likelihood. These initial zero-shot classifications could then be used as examples in a few-shot learning setup, gradually refining accuracy and improving results. Another promising direction is the integration of SpikeAgent with other specialized AI systems to create more comprehensive neuroscience research tools. For instance, combining SpikeAgent with agents specialized in behavioral analysis or multi-omics data interpretation could enable more holistic interpretations of neural data. Such multi-agent collaborations could tackle complex research questions by analyzing diverse data types simultaneously, potentially uncovering patterns and insights that might be missed by single-modality approaches. These advancements could significantly accelerate progress in areas such as brain-computer interfaces, computational neuroscience, and our understanding of neural dynamics in both health and disease.

In conclusion, SpikeAgent introduces a completely unexplored paradigm towards more explainable, scalable, efficient, and standardized neural data analysis. By combining LLMs with specialized visual processing capabilities, we have created a system that automates complex workflows while providing interpretable results. We envision, as this technology evolves, it has the potential to transform neuroscience research, introducing new ways to interact with neural data and potentially obtaining new or complementary insights into brain function and dysfunction and accelerating our understanding of neural dynamics.

## Methods

### SpikeAgent framework

We built SpikeAgent on LangChain and LangGraph^33^, frameworks designed to facilitate the development of language model applications by providing tools for prompts, document loaders, agents, memory, chat functionality, and more. By leveraging both LangChain and LangGraph frameworks, we established a robust foundation for developing stateful applications with LLMs. LangGraph allowed us to design cycle-capable workflows, which were critical for addressing the iterative nature of spike sorting tasks. We utilized multiple state-of-the-art LLMs in our system, including GPT-4, Claude-3, and Gemini, to leverage their zero-shot reasoning capabilities. We used the LLMs without fine-tuning to evaluate their effectiveness in handling specialized neuroscience tasks.

We implemented key features such as loops and conditionals in our spike sorting pipeline, automatic state-saving after each step, and human-in-the-loop capabilities. These frameworks enabled us to integrate various LLM providers, apply prompt engineering techniques, and incorporate external tools, allowing us to create a versatile and powerful spike sorting system with fine-grained control over both the flow and state of the application. We designed the SpikeAgent architecture to follow a ReAct (Reasoning and Acting) approach^34^, which we incorporated into the LLM’s tool-calling capabilities.

### Tools and modules

We integrated several specialized tools and modules into our SpikeAgent, leveraging a CodeAct^24^-inspired approach to consolidate actions into a unified, flexible code action space. This approach allowed us to perform dynamic code generation and execution, enhancing the system’s adaptability to various spike sorting scenarios and user inputs. We designed the core of our system to include tools for environment setup, data reading and preprocessing, spike sorting execution, visualization, and curation. Each tool generated relevant Python code tailored to the specific requirements of the task at hand. We executed this code using a Python REPL tool, enabling real-time adaptation and interaction with the data and results.

### Frontend

We built the user interface for SpikeAgent using the Streamlit Python package^35^, providing a comprehensive and intuitive platform for user interaction. We designed the main chat interface to support text-based queries and display the conversation history, including multimodal outputs such as text, images, and visualizations from the spike sorting process. We incorporated a sidebar with tabbed sections to enable conversation management (e.g., saving, loading, and summarizing histories), voice interaction (audio input and text-to-speech output), and image uploading.

We leveraged Streamlit’s session state for efficient state management across interactions and seamlessly integrated the frontend with the LangGraph-based backend. This design supported multimodal input, including text, voice, and images, as well as multimodal output, addressing diverse user preferences and research needs.

### Vision-language model capabilities

We implemented the VLM module using a dynamic prompt generation function that constructed tailored system prompts for different input modalities. This function adapted the prompt based on the presence of ISI data, quality metrics, and example spike IDs. We developed four prompt variants to handle waveform-only inputs, waveform with ISI distribution, and waveform with different pre-rejection strategies. We utilized multiple instances of state-of-the-art visual language models (e.g., GPT-4o, Claude-3.5-sonnet) as independent reviewers. Each reviewer processed the generated prompt alongside visual data, including waveform images and, when applicable, ISI distributions. We implemented an asynchronous processing pipeline to evaluate multiple spikes concurrently, improving overall efficiency.

The reviewers generated structured responses that included detailed reasoning, a spike score (0-1), and a classification (Good/Noise). We developed a voting mechanism to aggregate results from multiple reviewers, which mitigated individual model biases. For the pre-rejection approaches, we incorporated two preprocessing strategies to filter spikes based on predefined SNR and ISI violation thresholds: (1) a code-based approach, where exact values were used in if-else conditions to reject noise; and (2) a text-based approach, where threshold values were provided as text input, allowing SpikeAgent to incorporate them into its contextual reasoning for more advanced curation decisions.

We designed the module with modularity and extensibility in mind, enabling the seamless integration of new models or evaluation criteria. The output was structured to integrate smoothly with the broader SpikeAgent framework, providing both human-readable explanations and machine-processable classifications for downstream analysis. This parallelization accelerates the process while maintaining the detailed chain-of-thought reasoning provided by each reviewer.

### Spike sorting functions and frameworks

We developed an automated spike sorting agent that leveraged SpikeInterface^36^ as its core framework to seamlessly integrate various spike sorting algorithms, datasets, and preprocessing workflows. We relied on SpikeInterface’s modular design to unify diverse data formats, enable standardized preprocessing, and facilitate quality control.

We designed the agent to leverage SpikeInterface’s modular framework for handling data, preprocessing, spike sorting, postprocessing, and quality assessment. While we allowed users to instruct the agent to deviate from the standard pipeline for specific needs, we implemented a default workflow that began with data loading via the extractor’s module, ensuring compatibility with diverse formats such as Neuropixels and flexible neural probes. We incorporated preprocessing steps, such as bandpass filtering and common average referencing, through the preprocessing module to standardize inputs for analysis. The spike sorting itself was performed using the sorters module, which wrapped multiple algorithms and supported dynamic parameter tuning by the agent.

We utilized the postprocessing module to compute templates and unit locations and calculated quality metrics such as signal-to-noise ratio and isolation distance using the quality metrics module. We also integrated visualizations supported by the widget’s module, enabling the agent to plot relevant outputs for users. These metrics and visualizations were incorporated into the agent’s vision-language understanding capability, which used them to perform reasoned manual curation.

### Benchmarking of different LLM for autonomous spike sorting

The benchmarking of spike sorting agents was conducted using 13 LLMs, including OpenAI models (GPT-4o, GPT-4-turbo, GPT-o1, GPT-4o Mini, GPT-3.5 Turbo), Anthropic models (Claude-3.5-Sonnet, Claude-3-Opus, Claude-3-Haiku, Claude-3-Sonnet), and Gemini models (Gemini-1.5-Flash, Gemini-1.5-Flash-8b, Gemini-1.5-Pro, Gemini-2.0-Flash-Exp). The benchmarking was conducted to evaluate the models’ ability to generate consecutive code of spike sorting and tool chaining capability based on a predefined system prompt detailing the sequences of steps and the default parameters required for each step. The same initial prompt was provided to all models, forcing them to proceed through the end-to-end process. Each model underwent 100 iterations with different seeds, and data was collected on the number of steps completed before failing and the input/output token counts for each iteration.

Spike sorting was performed on flexible neural probe data using the Mountainsort4 algorithm. As this benchmarking focused on testing the code generation continuity of LLM models, SpikeInterface operations like spike sorting and extracting waveform were instead conducted by loading pre-saved sorting results and waveforms. The system prompts were originally designed to require user input for parameter decisions at each step; however, these constraints were removed for this benchmarking experiment to allow uninterrupted progression through the steps.

### Benchmarking SpikeAgent’ spike curation agreement with human expert

To benchmark the SpikeAgent’s curation agreement with human experts, we evaluated four VLMs capable of processing visual inputs: GPT-4o, GPT-4 Turbo, Claude 3.5 Sonnet, and Claude 3 Sonnet. These models were designed to process structured input sequences, beginning with a system prompt that provided task-specific instructions, examples of good and bad spike units, and visual information, such as waveforms and ISI distribution graphs. Using this input, the VLMs generated evaluations for each spike unit, assigning a score and providing a detailed rationale for their classification.

Four input formats were tested to evaluate model performance under varying conditions: (1) the average waveform from the peak amplitude channel, centered on a fixed 3 ms temporal window (1.0 ms pre-spike to 2.0 ms post-spike); (2) the waveform paired with an ISI distribution histogram displaying the 1.5 ms biophysical refractory period (x-axis range: 0–10 ms); (3) pre-filtered units excluding non-neuronal activity using signal quality thresholds; and (4) a multimodal combination of waveform images, ISI plots, and text-based SNR/ISI values derived from SpikeInterface computations.

We explicitly defined in the system prompt used in all configurations the features of high-quality spikes, including characteristics such as amplitude, peak clarity, noise level, and waveform reproducibility. For cases that included ISI distribution graphs, the prompt emphasized that ISI values should not fall below the biophysical refractory period of 1.5 ms. Based on these criteria, the VLMs assigned a normalized score (0 to 1) to each spike and provided a detailed explanation of their reasoning. To encourage interpretive flexibility and account for ambiguity in spike sorting, we set the model’s temperature parameter to 0.7.

### Time efficiency

We assessed the efficiency of VLM-based curation by comparing its performance to that of human curators. We measured processing speeds for VLMs using GPT-4o under three distinct operational modes. In the first mode, referred to as VLM asynchronous (VLMa), we utilized a maximum of 120 concurrent processes, allowing three reviewers to evaluate each spike simultaneously. This configuration enabled the simultaneous review of batches of 40 spikes. In the second mode, VLM3, we conducted synchronous reviews, where three reviewers collaboratively evaluated a single spike at a time. In the third mode, VLM1, a single reviewer processed all spikes sequentially, one by one. Each trial was repeated three times to calculate the average processing time and its variability, measured as the standard deviation.

To benchmark time efficiency of human curators, we had three human curators (one expert and two novices) evaluate spikes using waveform and ISI distribution images in the Neuropixels dataset. The curator assigned scores on a 0-to-1 scale across the four categories: amplitude, noise, shape, and reproducibility. We conducted a trial on datasets containing 40, 60, 80, and 100 potential spikes, recording the total execution time for each condition.

### Electrophysiology data

We used two distinct raw recording datasets obtained from two different mice: one recorded with a rigid probe and the other with a flexible probe.

The first dataset consisted of raw data collected using Neuropixels 2.0 probes, which are publicly available^5^. Briefly, the Neuropixels 2.0 probes feature a high-density array of recording sites, and an improved signal-to-noise ratio compared to earlier versions. In one of the recordings, we segmented the raw recordings into 1-minute snippets for analysis to standardize processing time and facilitate comparisons across experiments. The details of the probe geometry, experimental setup, and animal handling are thoroughly documented in the original dataset’s publication.

The second dataset was derived from 1 day of recording from flexible neural probe^37^. In short, the raw electrophysiological data were recorded using an Intan Technologies RHD2132 amplifier chip interfaced with a custom printed circuit board (PCB) for signal acquisition. Recordings were made at a 10 kHz sampling rate using a tetrode-like flexible electrode array, designed to maintain stability and biocompatibility during chronic implantation. We restricted the analysis to a 2-minute segment extracted from a 20-minute continuous recording session to optimize computational time without compromising data integrity. The details of the probe geometry, experimental setup, and animal handling are also documented and openly accessible in the original dataset’s publication.

## Data availability

Data used in this study will be made public with the manuscript publication.

## Code availability

The software code for this study will be made publicly available and maintained at the time of publication at http://github.com/LiuLab-Bioelectronics-Harvard/SpikeAgent.

All data analysis and visualization in this study were implemented using Python 3.10.13. The following packages and tools were used: LangGraph (v0.2.64), LangChain (v0.3.11), SpikeInterface (v0.101.1): https://github.com/SpikeInterface, Jupyter Notebook (v7.2.2), Seaborn (v0.13.2), Pandas (v2.2.1), SciPy (v1.14.1), NumPy (v1.26.4), Scikit-learn (v1.4.1.post1), Matplotlib (v3.9.3), UMAP-learn (v0.5.6), HDBSCAN (v0.8.40), and Streamlit (v1.41.1). Spike sorters: MountainSort 4 (v1.0.6): https://github.com/flatironinstitute/mountainsort, MountainSort 5 (v0.3.3): https://github.com/flatironinstitute/mountainsort, KiloSort 4 (v4.0.21): https://github.com/MouseLand/Kilosort, HerdingSpikes (v0.4.6): https://github.com/mhhennig/hs2, Tridesclous (v1.6.8): https://github.com/tridesclous/tridesclous

## Extended data figures and figure captions

**Extended Data Fig. 1:**
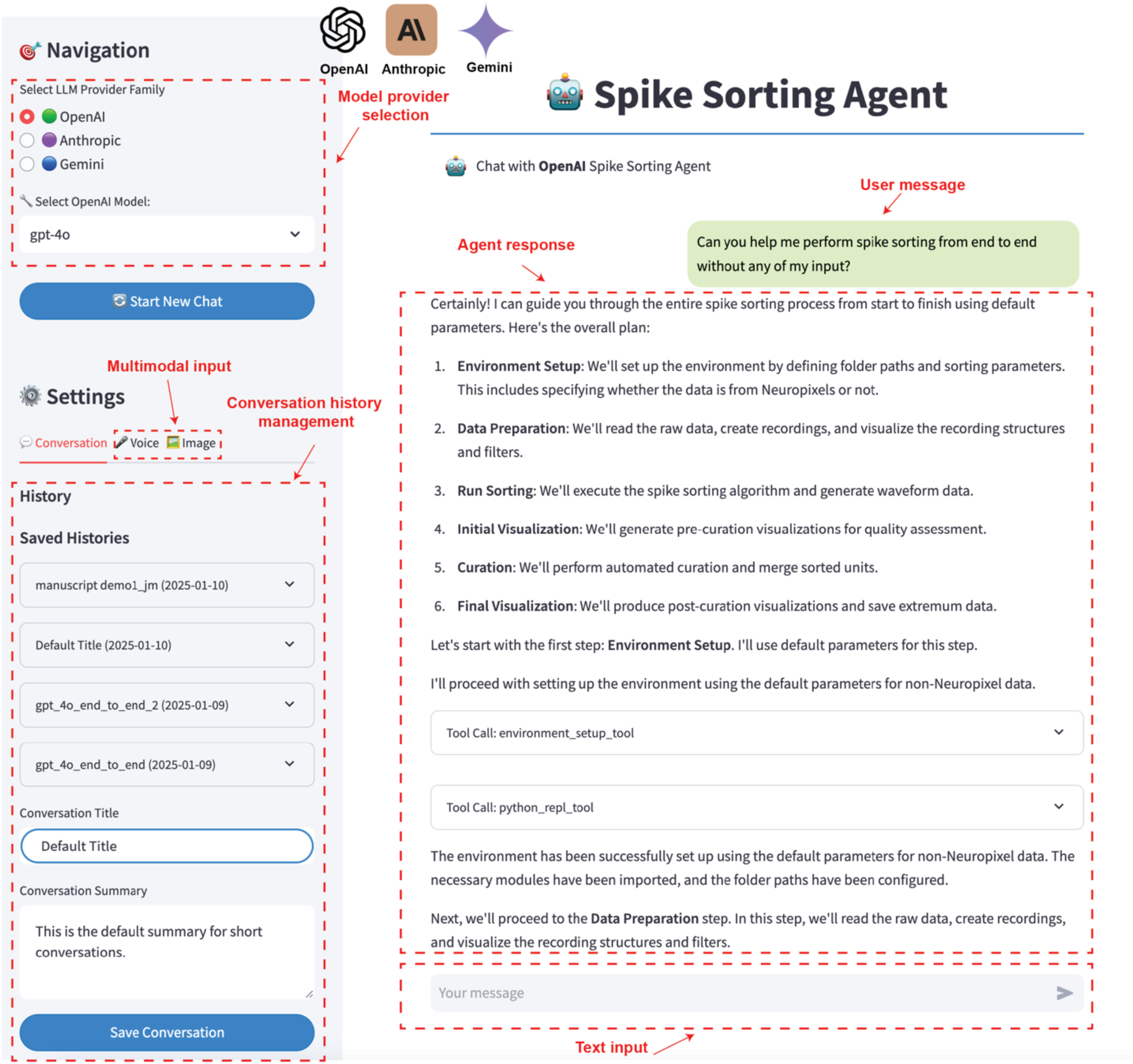
Frontend interface of the SpikeAgent. The interface provides an intuitive platform for performing spike sorting with multimodal input and model customization. The Navigation panel (left) allows users to select the desired model provider (e.g., OpenAI) and the specific model variant for spike sorting tasks. The Settings section offers multimodal input options, including text, voice, and image inputs, enhancing flexibility for various use cases. Users can manage conversation history through the Saved Histories feature, enabling easy access to previous sessions and seamless continuation of workflows. In the Agent response panel (center), the SpikeAgent interacts with the user by providing detailed step-by-step guidance for the end-to-end spike sorting pipeline, including environment setup, data preparation, sorting, curation, and final visualization. The outlined workflow ensures transparency and modularity in the spike sorting process. Users can initiate tasks with minimal input by providing a single text-based query in the Text input field (bottom right). The displayed conversation demonstrates the agent’s ability to handle queries autonomously, generating responses and executing the necessary tools to complete the task.

**Extended Data Fig. 2:**
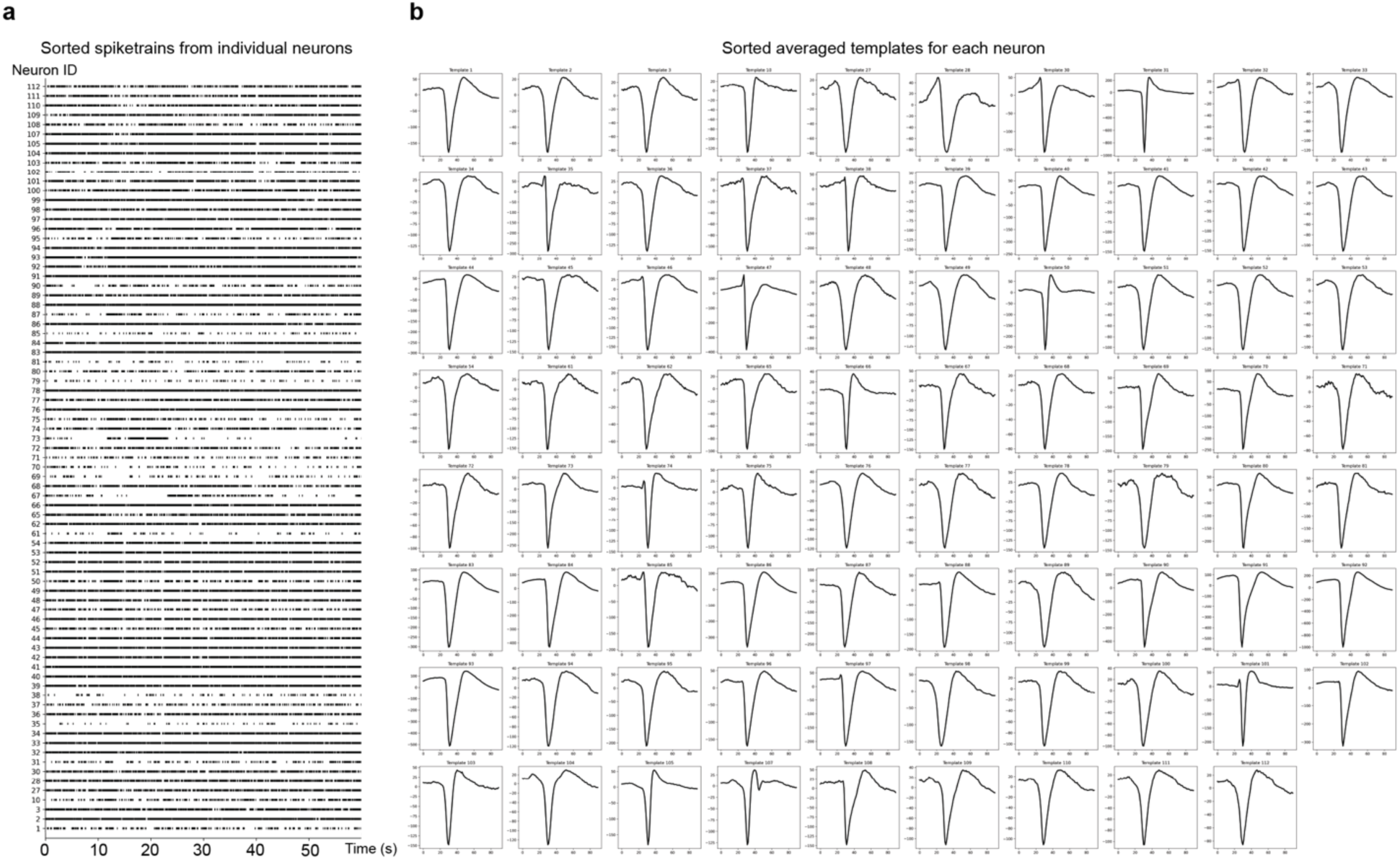
Spike sorting results from representative Neuropixels recording. **(a)** Sorted spike trains from individual neurons, labeled by unique Neuron IDs. Spike times are plotted over a 60-second recording interval. **(b)** Averaged spike templates for each neuron, aligned to peak amplitude and sorted across the recording duration. Templates represent mean waveforms derived from spike events detected over the full 60-second period

**Extended Data Fig. 3:**
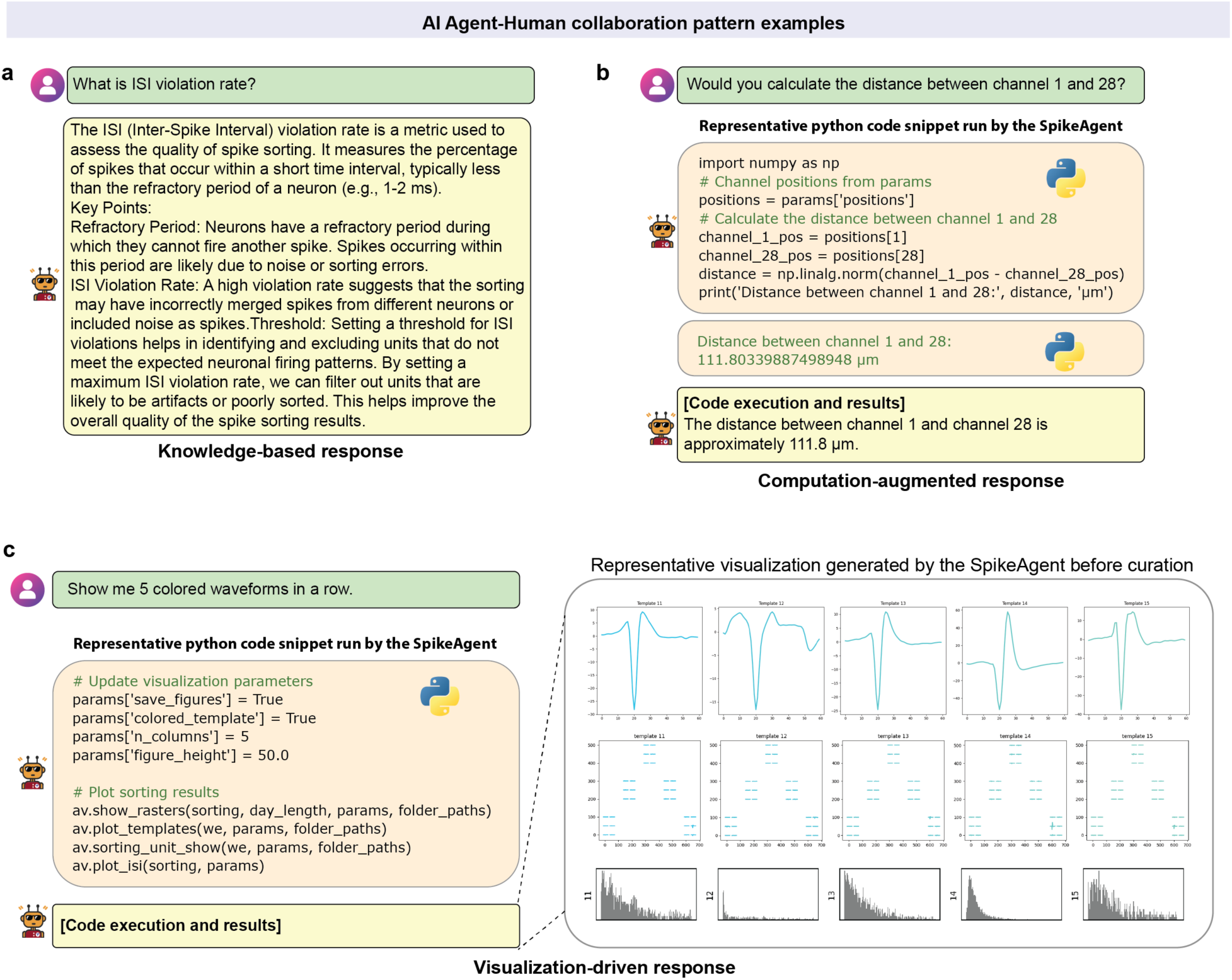
Example AI agent-human interaction patterns. **(a)** Knowledge-based responses: The SpikeAgent provides direct, detailed explanations to user queries by leveraging its extensive domain knowledge, without requiring additional computations or data processing. As an example, when asked about the ISI violation rate, the agent offers a comprehensive explanation of its definition, significance, and implications for spike sorting quality. **(b)** Computation-augmented responses: The SpikeAgent generates and executes code to perform calculations or data analysis, then interprets and presents the results in a user-friendly manner, combining its programming capabilities with domain expertise. As an example, when asked to calculate the distance between two channels, the agent writes Python code to compute the distance and presents the result with appropriate context. **(c)** Visualization-driven responses: The SpikeAgent creates complex, task-specific visualizations on demand by generating and executing appropriate code, translating user requests into informative graphical outputs. As an example, when asked to show 5 colored waveforms in a row, the agent creates a multi-panel figure displaying waveforms, templates, and ISI distributions, demonstrating its ability to produce sophisticated visualizations tailored to user needs.

**Extended Data Fig. 4:**
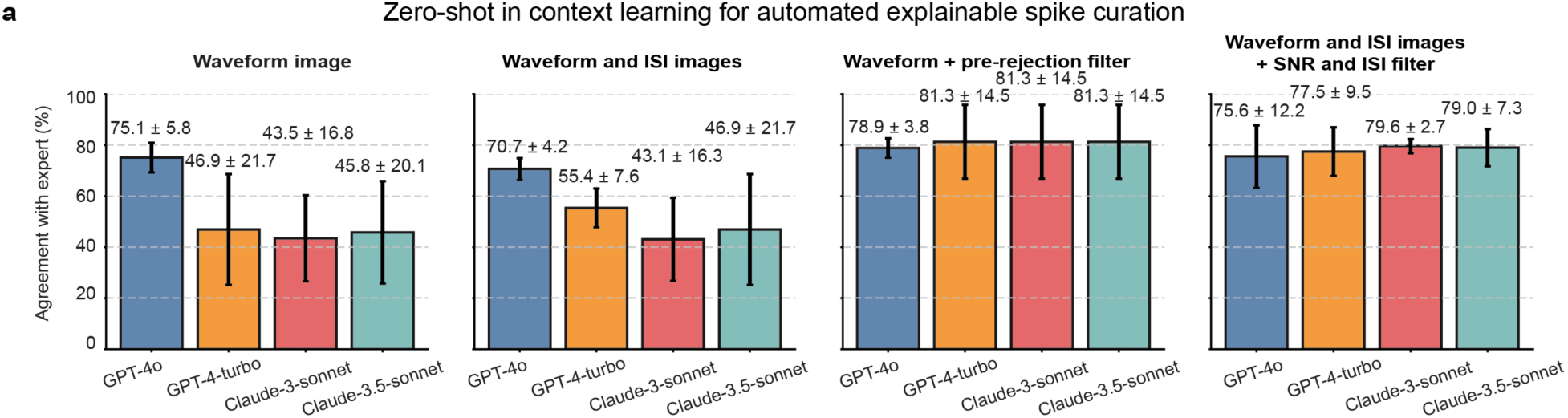
Zero-shot VLMs curation. **(a)** Agreement with human expert consensus comparison of VLM input modalities for spike curation in zero-shot in context learning. Bar plots show model performance under four conditions: waveform images alone, waveform with ISI images, waveform with ISI images after pre-rejection filtering, and waveform with ISI images plus text-based values of the SNR and ISI metrics. Data are mean ± s.d.

## Acknowledgements

We acknowledge all the Liu Lab members for their invaluable assistance with manual curation consensus and support to this work. A.ML. acknowledges the support from the RCC-Fellowship of Harvard University and the Excellence Fellowship of the Fundacion Rafael del Pino. We acknowledge support from NIH/NIDDK 1DP1DK130673 (J.Liu); NSF ECCS-2038603 (J.Liu and N.L.); and NIH/NLM 5R01LM014465 (J.Liu, N.L., and J.D.).

## Author contributions

Z.L., A.ML. and J.L. conceived the idea. Z.L., A.ML. and J.B. developed the method and performed experiments. Z.L. and A.ML. prepared figures and drafted the manuscript. All authors contributed critical discussions and input on the figures and results. J.L. supervised the study.

